# Methemoglobin aggregation is modulated by the anti-sickling drug voxelotor

**DOI:** 10.1101/2024.06.16.599216

**Authors:** Brandon Cove, Aldo Munoz, Arnav Singh, Gavin Jann, Kristal Brandon, Li Xing, Melanie J. Cocco

## Abstract

Sickle cell disease is caused by a mutation in the beta subunit of hemoglobin (HbSS) that drives Hb fiber formation when the protein is in the deoxygenated (tense, T) state. The drug voxelotor was recently approved to treat sickle cell disease by preventing HbSS fiber formation. Voxelotor acts as an allosteric inhibitor of polymerization by maintaining the HbSS protein in the relaxed (R) conformation, limiting polymerization of T-state fibers. Normal blood cells contain small amounts of natural Hb fibers and a few percent of the Fe^3+^ ferric form, metHb, incapable of binding oxygen. Although the drug Voxelotor is now in use, the effect of the drug on the oxidized metHb state has not been reported. Here we assessed the influence of voxelotor on normal human metHb. We compared the aggregation of metHb at two pH values (5.5 and 7.1). MetHb is known to form organized fiber structures at or below pH 5.5. We find that voxelotor significantly enhances fiber formation of metHb R-state at pH 5.5, consistent with the mode of action for this drug in maintaining the Hb R conformation. The opposite effect is observed at physiological pH values. Voxelotor significantly decreases the rate of metHb aggregate formation at pH 7.1 but did not affect protein stability. Notably, drug binding drives metHb into novel spherical particles with a morphology never seen before for Hb. The formation of these particles should be considered in patients being treated for sickle cell disease with voxelotor.

**WHY IT MATTERS:** Voxelotor is an FDA-approved drug for sickle cell anemia, known to prevent hemoglobin fiber formation. Here, we investigate its effect on methemoglobin, the form of hemoglobin in which iron takes on the ferric Fe^3+^ state. Our study examines voxelotor’s impact on methemoglobin aggregation and stability. At pH 7.1, we found voxelotor to have an effect on methemoglobin solubility as evidenced by the formation of novel methemoglobin spherical structures. We observe that voxelotor significantly increases methemoglobin fiber formation at pH 5.5 but, notably, reduces methemoglobin aggregation at physiological pH levels. Minimal impact on methemoglobin thermodynamic stability is noted. These findings suggest voxelotor’s potential therapeutic efficacy for various hemoglobinopathies, including conditions characterized by Heinz body formation.

## INTRODUCTION

Normal red blood cells (RBCs) contain a small amount of hemoglobin (Hb) in the oxidized Fe^3+^ ferric form, metHb, that is unable to bind oxygen. RBCs maintain a system of enzymes including Cyt b^5^ to reduce metHb to the functional Fe^2+^ state (oxy/deoxy Hb). When enzyme reduction systems operate most efficiently, intracellular metHb concentrations are ∼1%, but can be found up to levels of 6% in normal humans [1]. Methemoglobinemia occurs when levels of metHb are elevated. This can arise from genetic defects or from oxidizing agents like the analgesic benzocaine. Most individuals have no symptoms below an intracellular concentration of 30% metHb but show evidence of cyanosis above that level; metHb concentrations above 50% can be lethal [2]. The heme cofactor is released more readily in metHb; enhanced dissociation of subunits and protein unfolding make the metHb form less stable than the oxy/deoxy forms. Over time, metHb and denatured Hb aggregate to form large particles called Heinz bodies that can embed in the RBC membrane. These metHb aggregates affect cell membrane flexibility; their accumulation is a determining factor in RBC senescence [2].

Another pathology of hemoglobin aggregation is found in sickle cell disease, a genetic condition arising in individuals with a homozygous mutation of Glu6 to Val in the human β globin subunit (HbSS). The Glu6Val mutation stabilizes protein/protein interactions that drive HbSS fiber formation when the protein is in the tense (T), deoxygenated state. Fibers do not form when the protein is in the relaxed (R) oxy state. In the brief period after HbSS RBCs have off-loaded oxygen cargo, deoxy HbSS starts to polymerize, and sickled cells are formed. These HbSS sickled cells lose the flexibility required to passage through capillaries, obstruct blood flow and have a shortened cellular lifetime. The consequences for homozygous individuals (HbSS) are devastating including anemia, intense pain, stroke, organ failure, and death; collectively these are all part of sickle cell disease. **Figure 1A** shows a model of the HbSS fiber based on packing found in the crystal structure of deoxy HbSS [3].

**Figure 1.**
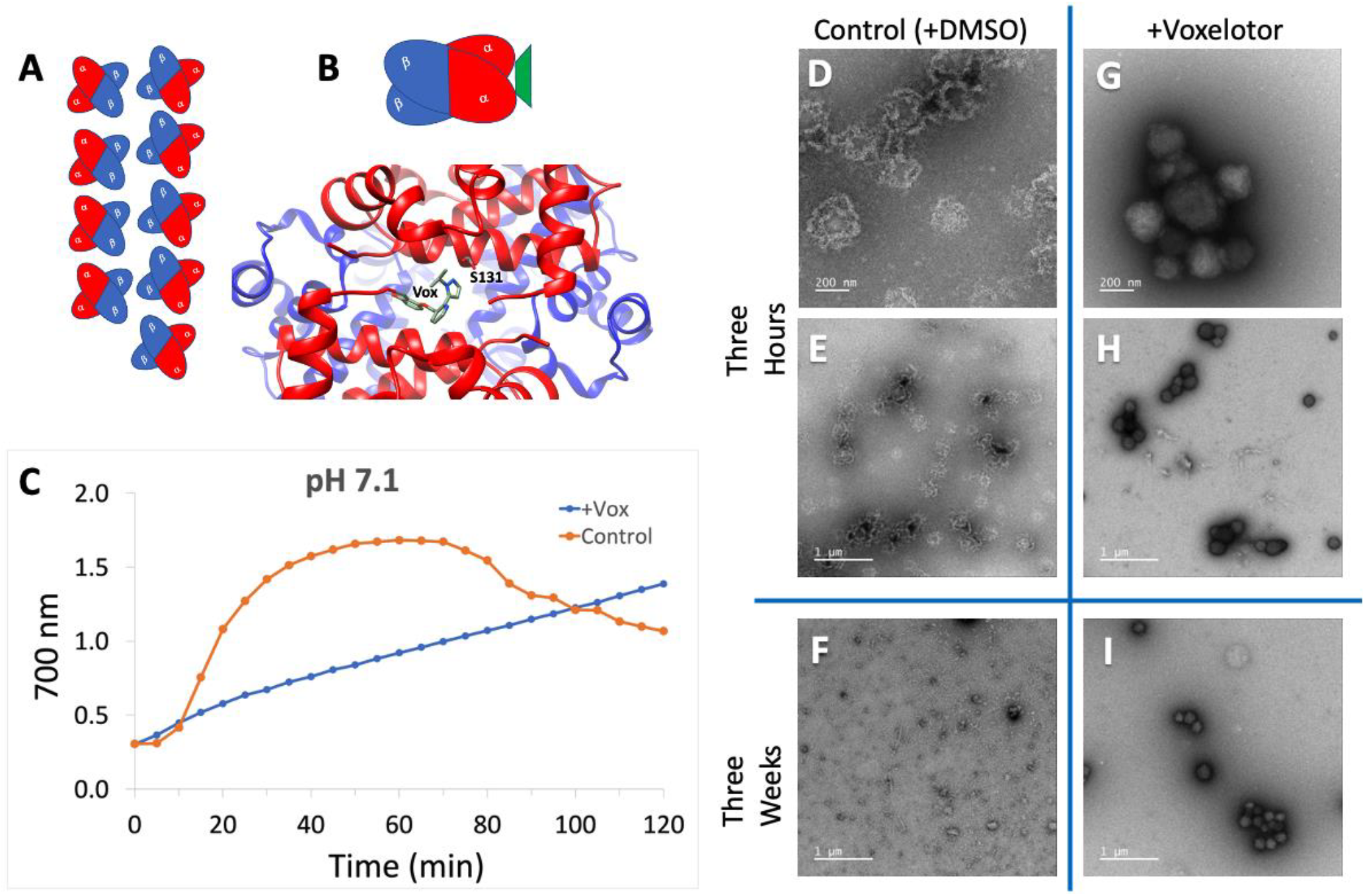
Particle formation at pH 7.1. **(A)** Model of deoxy HbSS fibers in the T state adapted from figure 8 in [3]. **(B)** The structure of voxelotor bound between HbSS α subunits (5E83.pdb). Voxelotor forms a covalent Schiff base with the N-terminus of one α subunit and hydrogen bonds to Ser131 on the other subunit, favoring the R state for the tetramer (model shown) to prevent fiber formation of HbSS. **C)** Effect of voxelotor on metHb particle formation measured by scattering at 700 nm, pH 7.1, 200 mM sodium phosphate, 37 °C; blue includes drug; orange is control. **D-I)** TEM, Effect of voxelotor at three hours at 37 °C, pH 7.1 and after 3 weeks at 4 °C. Control: **D)** three hours (200 nm scale); **E)** three hours (1 μm); **F)** three weeks (1 μm). With drug: **G)** three hours (200 nm); **H)** three hours (1 μm); **I)** three weeks (1 μm).

The drug voxelotor (GBT440 or Oxbryta) was recently approved to treat sickle cell disease by preventing HbSS fiber formation [4, 5]. Voxelotor forms a covalent Schiff base with the N-terminus of one Hb α subunit. Hydrogen bonding of the drug to Ser131 of the second α subunit maintains the Hb tetramer in the R conformation, even in the absence of oxygen ligand (**Figure 1B**) [5]. In their study of the voxelotor effectiveness, Oksenberg and colleagues found that deoxy HbSS incubated with drug had a slower rate of fiber particle formation determined by turbidity at 700 nm compared to protein without drug (Figure 2 in [5]). Since the deoxy T-conformation drives fiber formation of the sickle cell mutant, drug stabilization of the R-conformation is believed to be the mechanism by which deoxy HbSS remains soluble.

**Figure 2.**
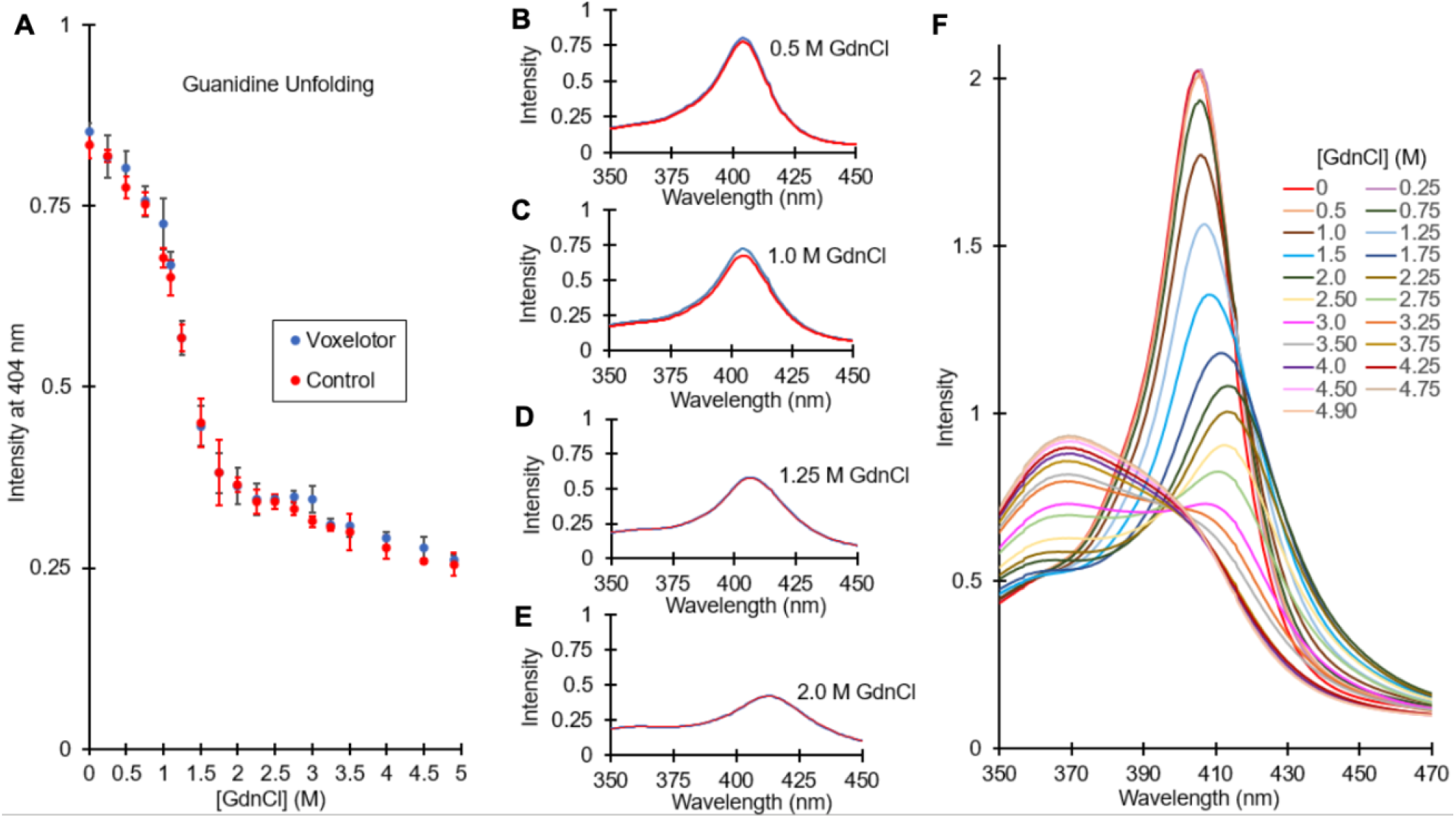
Stability of Hemoglobin. **(A)** Unfolding of the hemoglobin tetramer under varying concentrations of GdnCl (M) at a wavelength of 404 nm for both voxelotor (blue) and control (red, DMSO). **(B-E)** UV/Vis absorption intensity against wavelength (nm) for voxelotor (blue) compared to Control (red, DMSO) across different concentrations of GdnCl (0.5 M to 2.0 M) where hemichrome intermediate is populated. **(F)** UV/Vis absorption intensity against wavelength (nm) for voxelotor samples only (no control) across a range of GdnCl concentrations (0 M to 4.90 M).

Although many studies have assessed the influence of voxelotor on deoxy HbSS [4-6], oxidized metHb has not been studied with drug bound. Here we explore the effect of voxelotor binding on metHb at two pH values. We compare aggregation kinetics and the morphology of particles formed over time. In addition, we assessed the stability of the protein with voxelotor bound using chemical denaturation. Voxelotor has a significant influence on solubility and the protein/protein interactions that drive large aggregates but does not affect the thermodynamic stability of the protein.

## MATERIALS AND METHODS

### Materials

Human Methemoglobin (Ferrohemoglobin) was purchased from Sigma-Aldrich, Missouri, US (sigma-aldrich.com) Cat #H7379. Voxelotor (2-hydroxy-6 ((2-(1-isopropyl-1H-pyrazol-5-yl)pyridin-3-yl)methoxy) benzaldehyde) also known as GBT440 or trade name Oxbryta was purchased from MedKoo Biosciences, Inc, North Carolina, US (medkoo.com), Cat # 329516.

### Hb Aggregation Kinetics

A stock solution of 9.9 mM voxelotor was prepared in DMSO. 30 mg Hb were dissolved into 2 ml 0.2 M potassium phosphate buffer, pH = 7.1 to give a concentration of 233 μM Hb. Protein solutions were used immediately after preparation. MetHb samples were centrifuged and split into two 1-ml portions. 25 μl voxelotor stock was added to one sample (final drug concentration = 256 μM, 1.1 equivalents) and 25 μl DMSO to the other (control). We determined that 5 minutes was sufficient time for consistent kinetics and consider this time for the drug to fully react with Hb. The rate of particle formation with a 5 minute incubation time was identical to rates of longer incubation times. Voxelotor was allowed to incubate with Hb proteins for five minutes at room temperature, pH 7.1 and then absorbance (turbidity) was measured for up to two hours using a Beckmann DU800 spectrophotometer equipped with a Peltier temperature regulator set to 37 °C. For particle measurements at pH 5.5, the drug was first incubated with protein at pH 7 in H^2^O for an hour at 4 °C and then the pH was lowered with a pre-determined volume of acetic acid to achieve a final pH=5.5. The same procedure was applied to the control sample with 25 μl DMSO added.

### TEM Imaging

Protein samples were prepared under identical conditions as above. Samples were plated immediately at 3 hours incubation at 37 °C. Additional TEM plates were prepared from the same samples stored at 4 °C for three weeks. To prepare sample for TEM study, a droplet of the sample was applied on glow-discharged carbon-coated copper grids and negatively stained with 2% uranyl acetate solution. The sample grids were then examined with a JEM-2100F transmission electron microscope (JEOL, Japan) equipped with a field emission electron source and operated at 200 kV. Images were collected on a Ganta OneView camera (Ametek, USA) at magnification of 30,000x to illustrate the morphology of the sample.

### Chemical Denaturation

Guanidine hydrochloride (GdnCl) unfolding experiments were performed using methods described in [7]. 1.1 eq of voxelotor or DMSO were pipetted into one Hb sample; DMSO was added to the control and incubated at 4 °C for 5 minutes. Aliquots of protein solutions were transferred to conical centrifuge microtubes and diluted with an appropriate amount of GdnCl solution to a final metHb concentration of 30 μM. This concentration was chosen to ensure measurement of tetramer unfolding rather than the dimer [7]. Three separate samples were prepared for each GdnCl concentration using the same stock protein solution for both the control and voxelotor tubes. Absorbance spectra were collected from 325 nm to 700 nm for each GdnCl concentration (0 - 4.9 M), using 2 mm cuvettes to allow detection of the strong Soret peak.

## RESULTS

Ferric metHb is significantly less stable than either of the ferrous (reduced) oxy or deoxy forms. Consequently, metHb will start to denature within a short time under physiological ionic strength and temperature conditions. We tested the effect of voxelotor on the denaturation of metHb by measuring turbidity at 700 nm. **Figure 1C** shows an increase in scattering as a function of time indicating aggregation at 7.1, 37 ˚C. In each control experiment, we observed a decrease in turbidity about 80 minutes after the initial increase. We sealed and gently turned the cuvettes to mix the contents and found that the scattering signal returned. Consequently, the decrease in turbidity at long times arises from aggregates that fall to the bottom of the cuvette as they grow larger. To assess the influence of voxelotor on metHb aggregation, we mixed 1.1 equivalents of the drug with metHb and compared the aggregation to a control with DMSO alone added. Our results (**Figure 1C**) show that the drug does have an effect in stabilizing the soluble form of metHb with a marked decrease in the rate of particle formation. We used TEM to visualize the resulting particles formed by metHb alone or bound by voxelotor and found that the morphology changes dramatically when the drug is present. **Figure 1D-I** shows the results when metHb samples were incubated at 37 ˚C for three hours and then the same samples were stored at 4 ˚C for three weeks. In the control experiment, large metHb aggregates form rapidly and decompose with time. Notably, the presence of the drug drives metHb into a spherical morphology that has never been seen for any form of Hb. These 100-200 nm diameter spheres are highly soluble and remain stable for weeks. The collection of dye around the spheres often occurs when protein is packed tightly together, excluding the dye.

We assessed the stability of the hemoglobin fold to determine whether the changes in aggregate morphology of metHb at pH 7.1 were related to a surface property or the result of a change in the thermodynamic stability of protein when bound to drug. We measured the stability of the metHb using the chemical denaturant guanidine hydrochloride (GdnCl). The unfolding of metHb is known to be reversible in GdnCl and involves an intermediate set of species referred to as hemichrome [7]. **Figure 2A** shows the unfolding of metHb with or without voxelotor determined from the heme Soret (absorbance = 404 nm). The drug does not change the mid-point of unfolding and does not have an appreciable effect on the population of hemichrome intermediates (**Figure 2B-E**). An overlay of the full wavelength spectrum as a function of GdnCl concentration shows the same features as previously published for the protein alone (compare **Figure 2F** to Figure 3 in [7]). These measurements indicate that the drug has no effect on the stability of monodisperse metHb.

**Figure 3.**
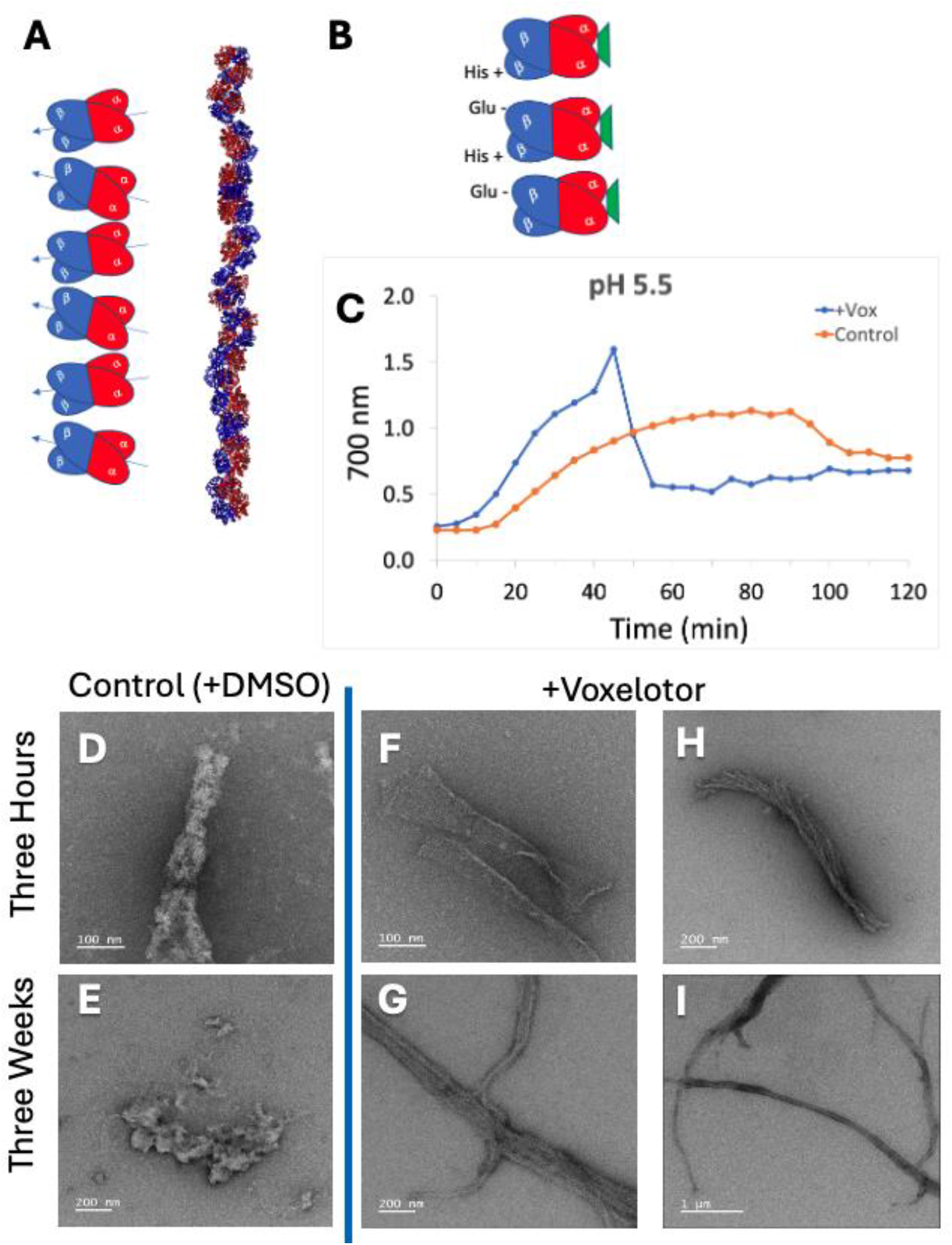
Particle formation at pH 5.5. **A)** Model and high-resolution crystal structure of metHb in the R-state at pH 5.5; one cell unit contains 12 tetramers (3ODQ.pdb). **B)** Model adapted from [8] showing the effect of voxelotor: both drug binding to promote the R-state and protonation of surface histidine at low pH drive fiber formation. **C)** Effect of voxelotor on metHb particle formation measured by scattering at 700 nm, pH 5.5, 200mM sodium phosphate, 37 °C; blue includes drug; orange is control. **D-I)** TEM, Effect of voxelotor at three hours at 37 °C, pH 5.5 and after 3 weeks at 4 °C. Control: **D)** three hours (100 nm scale); **E)** three weeks 200 nm scale. With drug: **F)** three hours 100 nm scale; **H)** three hours 200 nm scale; **G)** three weeks 200 nm scale; **I)** three weeks 1 μm scale.

Both oxyHb and metHb adopt the R conformation. Since voxelotor acts as an allosteric modulator to maintain the protein in the R conformation, we chose to study effect of the drug on metHb under conditions where the R conformation has been proven: fiber formation at low pH. MetHb will only form fibers at pH values below 5.5 [8]. To determine the structure of metHb, Larson, *et. al*., tested crystal growth at many pH values [8]. They found that the best crystals formed between pH = 4.5 to 5.5 and proposed that electrostatic interactions between β/β contacts would be optimal at low pH values and facilitate metHb polymer formation [8]. These fibers only form under conditions where histidine is positively charged. As voxelotor is believed to stabilize the R-state and this conformation is what makes up metHb fibers at low pH, the drug could be expected to increase particle formation at pH 5.5. Consequently, we tested the effect of voxelotor on the aggregation of metHb at pH = 5.5 (**Figure 3**).

**Figure 3A** includes the crystal structure of metHb fibers with 12 hemoglobin tetramers arranged linearly in a single unit cell [8]. Larson and colleagues proposed that two histidines can become protonated at pH 5.5 creating positively charged side chains that interact through favorable electrostatics with the negatively charged Glu6 of a neighboring Hb tetramer, driving protein/protein docking interactions (**Figure 3B** and figure 4 in [8]). MetHb fibers are distinct since the protein adopts the R-conformation in this single-strand fiber (**Figure 3A**). In contrast, HbSS fibers form at neutral pH with the protein the T-state (deoxy) conformation and HbSS fibers have a minimum requirement of two strands (**Figure 1A model**).

To study the protein with drug bound at pH 5.5, we had to optimize the chemistry of binding. The pK^a^ of the N-terminus of Hb is between 6.9-7.8 depending on the ligand bound to the heme; the N-terminus of Hb is not chemically reactive toward acylation below pH 6 [9]. We confirmed that the drug does not change turbidity measurements of metHb at pH 5.5 compared to control. Lack of drug binding would be expected at pH 5.5 since voxelotor requires formation of a Schiff base with the N-terminus of an α Hb subunit. The drug cannot react when the N-terminus is fully protonated. However, when the drug is incubated with metHb at pH 7 to allow covalent attachment to metHb, then the pH is lowered to 5.5, the drug is covalently bound and has a pronounced effect on metHb fiber formation (**Figure 3A**). Using the same turbidity measurement performed above at pH 7.1, we monitored particle formation with and without drug at pH 5.5 (after preincubation of the drug at pH 7). In contrast to the results at pH 7.1, the drug increases particle formation at the lower pH. TEM images shown in **Figure 3D-I** reveal a profound difference in the particle morphology. R-state fibers are indeed stabilized by the drug when the surface histidine side chains are protonated. Notably, the protein without drug does form fibers within three hours, but these decompose with time. The fibers formed with the drug bound are stable and quite long lived.

## DISCUSSION

We find that voxelotor slows the formation of denatured metHb aggregates at pH 7.1 to favor a spherical shaped particle that is stable for weeks. Voxelotor binding favors the formation of the R-state conformation for metHb in solution seen at low pH values. The drug stabilizes distinct particle morphologies at pH 7.1 and 5.5 because of different protein surface charge features. Positively charged histidines that lead to fiber formation at pH 5.5 are most likely the dominant force driving fibers even when voxelotor is bound. Although we do not yet have a high-resolution structure of the metHb spheres, we can say that the protein/protein interactions that produce spheres at pH 7.1 are not competitive with surface histidine/glutamate interactions at pH 5.5 since the spheres do not form at the lower pH. This is perhaps not surprising since electrostatic interactions are often among the strongest when considering protein structure.

Since we find that voxelotor does not change the stability of the hemoglobin fold, the formation of spherical particles at pH 7.1 is likely to be related to a surface property of the protein with drug bound. This is similar to the case of the sickle-cell mutant (HbSS) that is stable and functional but has a surface property that drives protein/protein aggregation and fiber formation. At pH 7.1, voxelotor drives metHb to form unique spherical particles that stabilize the protein and prevent aggregation of unfolded states.

Oxidized metHb and denatured Hb aggregate to form large Heinz bodies embedded or associated with the cell membrane. The formation of Heinz bodies plays a role in RBC senescence. Notably, voxelotor drives metHb into soluble spheres. Patients taking voxelotor to treat sickle cell disease may have altered metHb processing/outcomes since the drug stays covalently attached regardless of oxidation state. Evidence has revealed that oxidized RBCs are associated with microhemorrhages in the brain [10-12]. These microhemorrhages are a source of microbleeds that can cause both ischemic and hemorrhagic stroke among other severely damaging conditions (reviewed in [13]). It is possible that the formation of metHb-voxeolotor spheres might inhibit Heinz body formation. Studies are ongoing to determine if voxelotor could be repurposed to modulate Heinz bodies produced upon RBC aging.

## ACKNOWLEDGEMENTS

AM was supported by a MARC fellowship (NIH T34GM136498)

EM images from the UC Irvine Materials Research Institute (IMRI), supported in part by the NSF through the UC Irvine Materials Research Science and Engineering Center (DMR-2011967).

Global Blood Therapeutics (now part of Pfizer) has provided funding for the Cocco lab; however the studies described here were not part of that project.

## AUTHOR CONTRIBUTIONS

BC, AM, AS, GJ, KB, and LX performed experiments; AM and BC analyzed data and prepared figures. MJC designed the study and experimental protocol, analyzed data, and prepared figures. The manuscript was written through contributions of both first authors. All authors have given approval to the final version of the manuscript.

